# Jump-starting the T cell response in established tumors

**DOI:** 10.64898/2026.05.28.728503

**Authors:** Tyler R McCaw, Mingyong Liu, Christopher D Scharer, Jeremy M Boss, Tian Mi, Alexander F Rosenberg, Mei Li, Rebecca C Arend, Donald J Buchsbaum, James Markert, Troy D Randall

**Author notes:** Corresponding author: Troy D. Randall, Department of Medicine, Division of Clinical Immunology and Rheumatology University of Alabama at Birmingham, 1720 2nd AVE S, Birmingham, AL 35294.

## Abstract

Checkpoint blockade only works in 10-20% of patients. Consequently, investigators are testing checkpoint inhibitors in combination with drugs like the class I histone deacetylase inhibitor, entinostat (ENT). Unfortunately, the combination of ENT and checkpoint blockade fared poorly in patients with breast or ovarian cancer, despite promising pre-clinical results. Here we show that ENT enhances CD8^+^ T cell responses by maintaining a progenitor-like population of CD8^+^ T cells that supplies activated effector T cells to tumors for prolonged periods. Surprisingly, the anti-tumor effects of ENT are only experienced when delivered during a narrow window that occurs after T cell activation and before T cell exhaustion—a window that is likely closed in most patients. However, by first “jump-starting” the T cell response using an oncolytic virus, the anti-tumor activity of ENT and PD1 blockade is restored. These data establish a general paradigm, independent of tumor type, to rationally manipulate anti-tumor immunity.

## INTRODUCTION

Antibody-mediated blockade of inhibitory receptors on T cells is thought to “release the brakes” on anti-tumor immunity and eliminate tumors [1]. In fact, when successful, these therapies can trigger rapid tumor regression and prolonged remission [2]. Unfortunately, most patients with solid tumors do not respond to checkpoint inhibition [2,3]. Thus, we urgently need new ways of stimulating anti-tumor immunity and limiting off-target immune reactions. Histone deacetylase (HDAC) inhibitors were initially developed to impair tumor cell proliferation and promote terminal differentiation and apoptosis [4], effects due in part to increased histone acetylation, opened chromatin and altered gene expression [5]. For example, class I HDAC inhibitors increase tumor cell expression of major histocompatibility complex (MHC) proteins, costimulatory molecules, and inflammatory chemokines [6–8]. The activities of numerous non-histone proteins are also regulated by acetylation [9,10], including a variety of transcription factors, such as MYC, STAT3 and FOXP3, which regulate T cell function [11,12]. Thus, the anti-tumor effect of HDAC inhibitors is partly due to their effects on T cells [8,13–15].

Despite promising data in mice showing that HDAC inhibitors can enhance tumor-reactive T cell function and sensitize tumors to the effects of PD-1 blockade [16,15], these effects have not been replicated in the clinic [17]. In fact, the class I HDAC inhibitor, entinostat (ENT), recently failed to achieve study endpoints in two Phase 1b/2 clinical trials in which it was combined with PD1 blockade to treat recurrent epithelial ovarian cancer (NCT02915523) and triple negative breast cancer (NCT02708680). How can these results be reconciled? One possibility is that ENT is being administered without considering how or when it acts on T cells. In this regard, the epigenetic landscape of T cells is tightly linked to their activation status [18]. The majority of chromatin in resting, naïve T cells is closed and genes associated with T cell effector function are not expressed. After activation, T cells rapidly alter their chromatin accessibility, particularly at loci involved in proliferation and the expression of effector molecules. Many of these regions return to a closed configuration as activated T cells become exhausted or transition to memory [19,20]. Thus, the ability of HDAC inhibitors to promote anti-tumor immunity may be contingent upon the activation status of tumor-reactive T cells at the time the drug is administered [21].

Here we tested this hypothesis in a mouse model of breast cancer. We found that ENT suppressed the in vitro activation and proliferation of naïve CD8^+^ T cells, but did not prevent the proliferation or effector functions of previously activated T cells. In vivo, ENT potently enhanced anti-tumor CD8^+^ T cell responses by sustaining a progenitor-like population of CD8^+^ T cells that robustly generated tumor-reactive effector T cells for prolonged periods. Interestingly, the anti-tumor effects of ENT were only experienced when delivered during a narrow temporal window after tumor implantation, a time frame that corresponded with the peak of acute T cell activation. Importantly, ENT no longer enhanced anti-tumor immunity once T cells became dysfunctional, a result that likely explains the lackluster efficacy of ENT in human breast and ovarian cancer. However, by first “jump-starting” the T cell response to established tumors using an oncolytic virus, the anti-tumor activity of ENT and PD1 blockade was restored in mice with established tumors. These data establish a general paradigm, independent of tumor type, to rationally manipulate anti-tumor immunity and improve the efficacy of checkpoint blockade.

## RESULTS

### Timing of ENT administration dictates CD8^+^ T cell function and tumor control

To test whether ENT could impair T cell activation, we stimulated naïve, CTV-labeled CD8^+^ T cells in vitro with anti-CD3 and anti-CD28 in the presence of increasing concentrations of ENT. Low doses of ENT minimally impacted T cell proliferation, whereas higher doses abrogated proliferation (**Figure 1A**). However, increasing concentrations of anti-CD3 and anti-CD28 overcame the inhibitory effects of ENT and restored CD8^+^ T cell proliferation and PD1 expression (**Figure 1B-C**).

**Figure 1.**
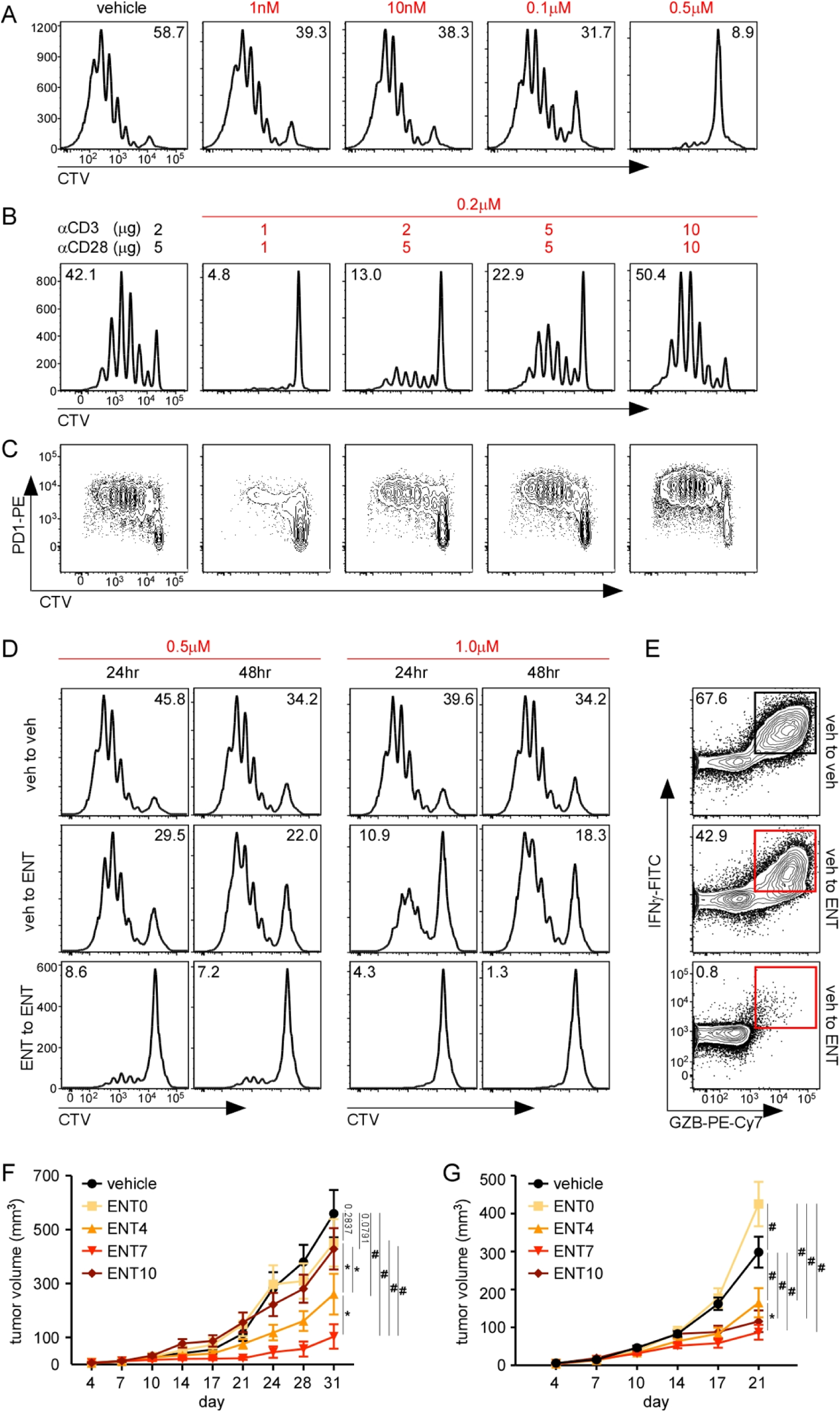
T cell activation state determines the impact of HDAC inhibition in vivo and in vitro. A-C. CD8 T cell proliferation measured by cell trace violet (CTV) dilution, n=2. **A.** CD8 T cells isolated from naïve mouse spleens activated using anti-CD3/CD28 in culture with escalating concentrations of ENT present from the start. **B-C.** Naïve CD8 T cell activated using increasing amounts of anti-CD3/CD28 in the presence of 0.2μM ENT. CTV dilution (**B**) and increasing PD-1 expression (**C**) are shown. **D.** CD8 T cells from naïve mouse spleens were activated using 2μg/5μg of anti-CD3/CD28 in the presence of vehicle or 0.5μM or 1.0μM ENT. Cells were transferred after 24 or 48 hours culture into fresh media containing vehicle or ENT and plate bound anti-CD3/CD28. All conditions harvested at 72 hours, n=2. CD8 T cell proliferation assessed by CTV dilution. **E.** Intracellular staining for IFNγ and GZB at the end of the 72 hour culture. **F-G.** Tumor growth curves of TS/A (**F**) and MC38-ova (**G**) tumors when ENT treatment is started at different days after tumor cell injection. ENT treatment started on day 0 (yellow), 4 (orange), 7 (red), or 10 (brown). Two independent experiments with n=6 per cohort per experiment. Significance determined using two-way ANOVA and Tukey’s correction for multiple comparisons. Mean and SEM shown. *p<0.05, **p<0.005, ^#^p<0.0005.

To test whether the inhibitory effects of ENT were restricted to the initial activation of CD8^+^ T cells, we stimulated naïve, CTV-labeled CD8^+^ T cells with anti-CD3 and anti-CD28 in media containing vehicle or ENT and transferred them into fresh media containing ENT or vehicle 24 or 48 hours later. T cells activated without ENT (veh to veh) proliferated robustly (**Figure 1D**), addition of ENT at either 24 or 48 hours only modestly reduced proliferation (veh to ENT), whereas the presence of ENT throughout culture (ENT to ENT) abrogated T cell proliferation (**Figure 1D**). Moreover, T cells activated without ENT expressed both IFNγ and granzyme B (GZMB), even if ENT was added at 24 hours, whereas T cells activated in the presence of ENT failed to express IFNγ or GZMB (**Figure 1E**).

To test whether the timing of HDAC inhibition altered its effects on anti-tumor immunity, we implanted mice with TS/A mammary adenocarcinoma cells and treated them daily with ENT or vehicle for 2 weeks beginning on days 0, 4, 7, or 10. Beginning on day 0 or 10 failed to slow tumor growth compared to vehicle controls (**Figure 1F**), whereas beginning treatment on day 4 moderately reduced tumor growth and beginning treatment on day 7 dramatically reduced tumor growth, with about a third of mice completely rejecting their tumors (**Figure 1F**). We obtained similar results using MC38-OVA tumors, demonstrating that the effect was not cell line or mouse strain-specific (**Figure 1G**). Importantly, the anti-tumor effects of ENT were dependent on CD8^+^ T cells for both TS/A and MC38-OVA tumors [15] (**Supplemental Figure 2**).

### ENT alters tumor-infiltrating CD8^+^ T cell activity

To determine how ENT affected tumor-infiltrating CD8^+^ T cells, we treated TS/A tumor-bearing mice with vehicle or ENT daily beginning at day 7 and performed RNAseq on CD8^+^ T cells isolated from tumors on day 10 and day 17. Principle component analysis revealed that CD8^+^ T cells from control and ENT-treated tumors clustered separately (**Figure 2A**). We found 3395 differentially expressed genes (DEGs) between control and ENT-treated samples on day 10, and 2546 DEGs between control and ENT-treated samples on day 17. Only 1017 DEGs were shared between day 10 and day 17 (**Figure 2B**). Many of the genes upregulated on day 10 were related to metabolic and effector pathways, whereas genes upregulated on day 17 were related to proliferation (**Figure 2C**).

**Figure 2.**
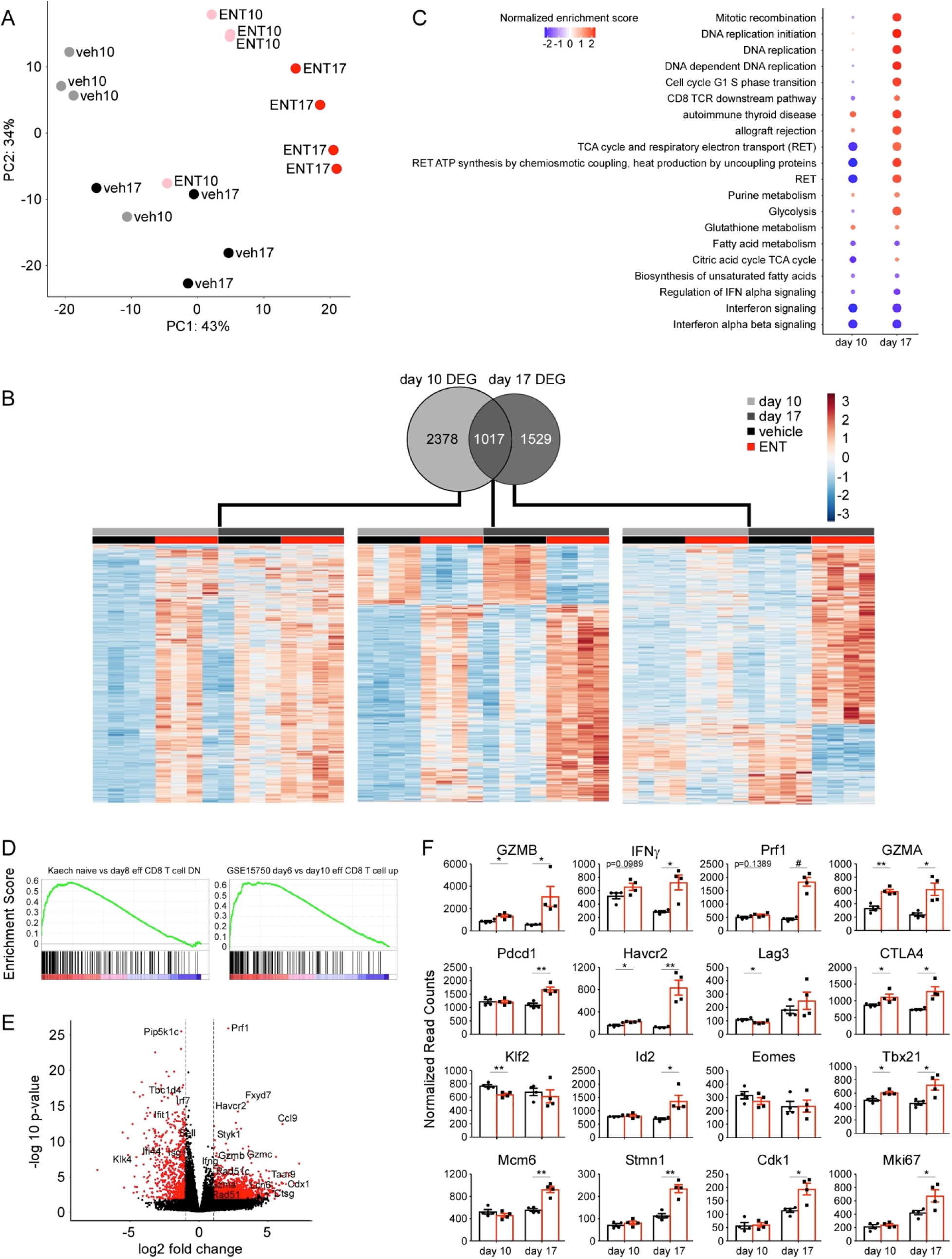
HDAC inhibition drives treatment duration-dependent changes in proliferation and effector function of tumor-infiltrating CD8 T cells. TS/A tumor-bearing mice were treated with ENT or vehicle starting on day 7 and tumor-infiltrating CD8 T cells sorted on day 10 or 17 for bulk RNA-sequencing, n=4 per cohort per time point. **A.** Principle component analysis of day 10 versus day 17 samples. **B.** Venn diagram and corresponding heatmaps of DEGs on day 10, day 17, or persistently differentially expressed (day 10 and day 17). **C.** Dot plot showing pathway analysis comparing day 10 ENT to vehicle and day 17 ENT to vehicle samples. Color indicates normalized enrichment score and circle size corresponds to –log10(p-value). **D.** GSEA of day 17 ENT versus vehicle samples comparing to specific datasets. **E.** Volcano plot of day 17 ENT versus vehicle showing most strongly DEGs. **F.** Normalized read counts of select genes, showing effector function, relevant transcription factors, T cell activation and exhaustion, and proliferation. Statistical significance determined by independent t-tests at each time. Mean and SEM shown. *p<0.05, **p<0.005, ^#^p<0.0005.

Pathway and gene set enrichment analysis (GSEA) showed positive enrichment of genes associated with CD8^+^ T cell effector function, DNA replication and oxidative phosphorylation (**Figure 2C-D**), but negative enrichment of genes linked to type I IFN signaling [22–24]. Accordingly, IFN-stimulated genes, including Ifi44, Ifit1 and IRF7, are highly enriched in vehicle-treated cells (**Figure 2E**), whereas T cell effector molecules like Prf1, Gzmb, Gzmc and Ccl9 were enriched in ENT-treated samples (**Figure 2E**). We also observed time-dependent increases in the expression of effector molecules like Gzmb, IFNγ and Prf1 in ENT-treated samples as well as increases in transcription factors like Tbet and Id2 (**Figure 2F**) [25,26]. In addition, the activation markers, PD1, TIM3 and CTLA4 were increased in ENT-treated samples, as were the proliferation-associated factors, Mcm6, Stmn1, Cdk1 and Mki67 (**Figure 2F**). Using flow cytometry, we confirmed that ENT treatment progressively increased the expression of IFNγ, GZMB and TNF (**Supplemental Figure 3A-D**). We also showed that ENT increased or sustained expression of the transcription factors, Tbet, Id2 and Tcf1 (**Supplemental Figure 3E-J**) [25–27]. Finally, we confirmed a higher frequency of Ki67-expressing CD8^+^ T cells in ENT treated tumors (**Supplemental Figure 3K-L**).

### ENT increases distinct populations of proliferating and effector CD8^+^ T cells

To test whether ENT altered the distribution of CD8^+^ T cell subsets, we sorted CD8^+^ T cells from ENT-treated or control tumors on day 17 and performed single cell RNAseq. Most cells from ENT-treated tumors clustered separately from those in control tumors (**Figure 3A**). Subclustering analysis revealed CD8^+^ T cells from ENT-treated tumors were enriched in clusters 1, 2, and 5, whereas CD8^+^ T cells from control tumors were enriched in clusters 0, 3, 4, 6, 7, and 10 (**Figure 3B-C**). Signatures of effector cells were enriched in clusters 1 and 2, whereas signatures of proliferating cells were enriched in cluster 5 (**Figure 3D**). By comparing the top 10 DEGs from each cluster (**Figure 3E**), we found that cluster 1 highly expressed numerous GZMs, implying that these cells were robust effectors. Similarly, cluster 2 highly expressed Ly6a, Ly6c, Galectin 3 and Plac8, genes associated with effector or effector-memory like cells. Impressively, cells in cluster 5 highly expressed proliferation-associated genes, including Stmn1, Tuba1b, Top2a and Cks1b (**Figure 3E**). Conversely, control cells in clusters 0, 3, 4 and 7 expressed higher amounts of the transcription factor, KLF2 (**Figure 3E**), which is primarily expressed in quiescent T cells and is downregulated in tissue resident CD8^+^ T cells [28]. Subcluster characteristics are cataloged in **Supplemental Table 1**.

**Figure 3.**
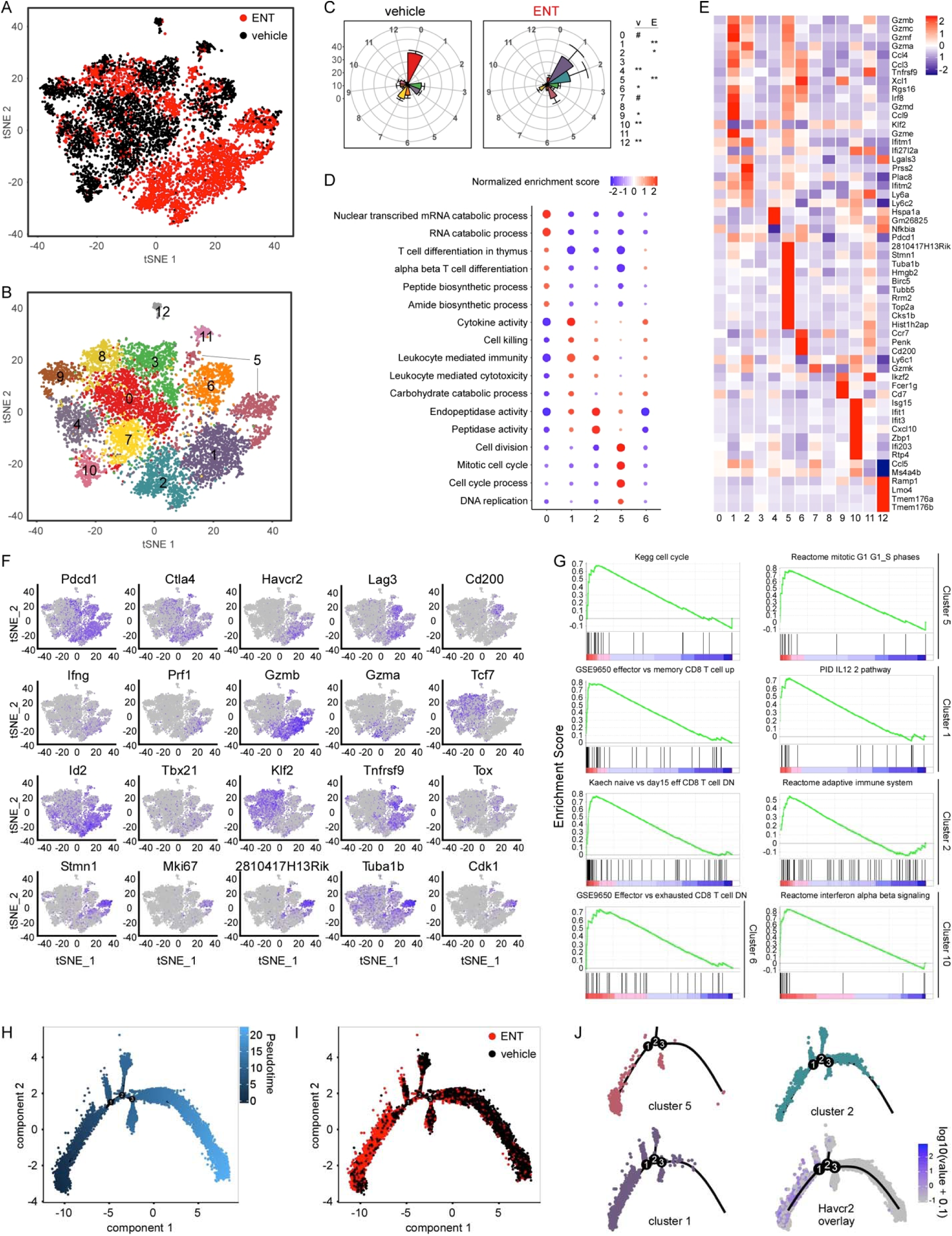
Single cell RNA-sequencing shows generation of highly proliferative and robust effector CD8 T cell populations in the tumor. TS/A tumor-bearing mice were treated with ENT or vehicle starting on day 7 and tumor-infiltrating CD8 T cells sorted on day 17 for single cell RNA-sequencing. Two independent experiments, n=2 per cohort per experiment. **A.** tSNE plot overlaid with sample treatment, vehicle (black) or ENT (red). **B.** tSNE plot with cluster analysis overlaid. **C.** Rose plot showing frequencies of vehicle and ENT treated cells in each cluster. **D.** Dot plot showing pathway analysis by cluster number (listed across bottom) where color indicates normalized enrichment score and circle size corresponds to –log10(p-value). **E.** Heatmap of top 10 DEGs in each cluster. Overlapping DEGs between clusters reduces the total number of genes shown. **F.** tSNE plots overlaid with select genes to show distribution of expression among clusters. Genes representing effector function, transcription factors, T cell activation and exhaustion, and proliferative capacity demonstrate concordance with bulk RNA-sequencing.**G.** GSEA of select clusters demonstrating their functional similarity to published data sets. **H.** Semi-supervised trajectory analysis using DEGs from initial analysis overlaid with psuedotime progression. **I.** Trajectory plot overlaid with treatment, vehicle (black) or ENT (red). **J.** Trajectory plot overlaid with cluster 5 (top left), cluster 1 (bottom left), or cluster 2 (top right), showing their respective increases in psuedotime or distance from the root. TIM3 expression overlaid (bottom right) shows it is largely found in the trunk.

We confirmed these results by overlaying the expression of individual genes on the tSNE plot (**Figure 3F**). Note that cluster 5 is spatially separated into two parts (**Figure 3B**) and that each possesses the proliferation signature (**Figure 3F**). Interestingly, cells in cluster 6, mostly from control tumors, strongly express multiple activation and inhibitory receptors, PD1, CTLA4, LAG3 and CD200, but notably lack TIM3 (**Figure 3E-F**). This phenotype is consistent with a dysfunctional profile, as co-expression of multiple inhibitory receptors, particularly LAG3 and CD200 family members, more reliably reflects exhaustion [29–31]. Finally, we used GSEA to confirm the proliferating signature of cluster 5 and the effector signature of clusters 1 and 2 (**Figure 3G**). Interestingly, cluster 6 and cluster 10, each predominantly comprised of cells from control tumors, had gene expression patterns associated with exhaustion and type I IFN signaling, respectively (**Figure 3G**).

We next conducted semi-supervised trajectory analysis, which arranged cells from left to right (low to high pseudotime) in an arc with only 3 small branches (**Figure 3H**). Most cells from ENT-treated tumors were placed on the left, close to the root of the trajectory, whereas most cells from control tumors were placed on the right (**Figure 3I**). By overlaying individual clusters on the trajectory map, we found that cluster 5 formed the root of the trajectory, cluster 1 started near the root and continued about halfway up the arc and that cluster 2 started near the root and continued over the top of the arc (**Figure 3J**). In contrast, clusters 0, 3, 4, 7, and 10, the major populations in control tumors, all mapped to the tail end of the trajectory (**Supplemental Figure 4**). Interestingly, cells that expressed TIM3 (Havcr2) were limited to the left side of the arc consistent with their position in clusters 1 and 2 (**Figure 3J**). These data suggest that ENT expands a population of proliferating, progenitor-like CD8^+^ T cells in cluster 5 that robustly generates effector cells in clusters 1 and 2.

### PD1^+^Tim3- progenitor T cells give rise to PD1^+^Tim3^+^ effector T cells

Based on gene expression (**Figure 3F**), the proliferating cells in cluster 5 should not express TIM3. In fact, we found a population of TIM3^-^Ki67^+^ cells in both ENT-treated and control tumors, which likely contains cells in cluster 5 (**Figure 4A-B**). We also found that ENT dramatically expanded a population of TIM3^+^Ki67^+^ cells (**Figure 4A-B**), which likely corresponds to effector cells in clusters 1 and 2. We observed similar results when we characterized tumor-specific T cells using a MuLV(env) MHC class I tetramer (**Supplemental Figure 5A**). Interestingly, the plots of TIM3 vs Ki67 are strikingly similar to the plots of TIM3 vs PD1 (**Figure 4A-D**). Moreover, we found that TIM3^-^PD1^+^ cells express high amounts of IL7Rα (CD127) and ICOS (**Figure 4E-F**), consistent with a memory-like state [32].

**Figure 4.**
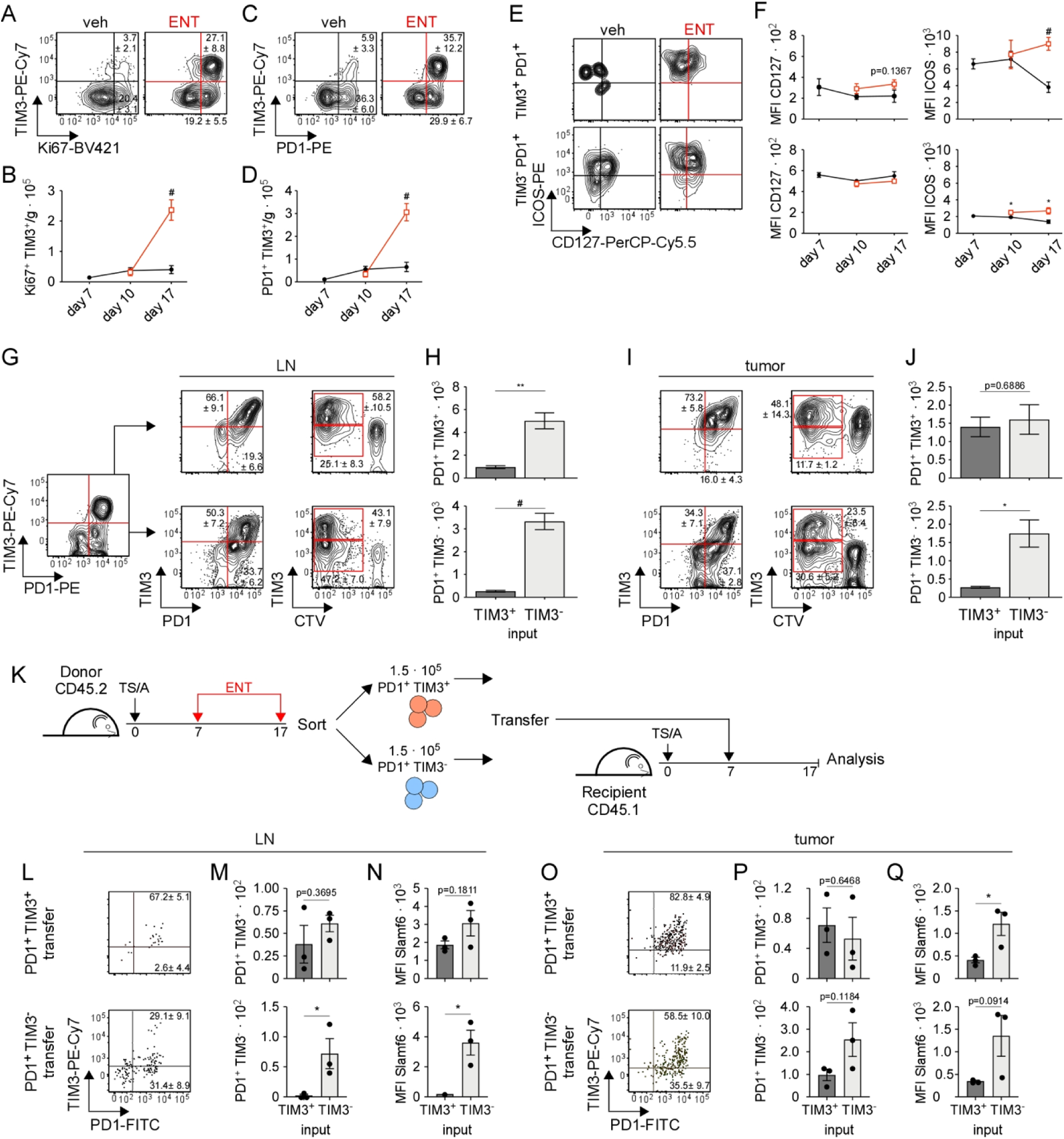
ENT-induced proliferative CD8 T cells are progenitors that give rise to effector CD8 T cell populations. A-F. TS/A tumor-bearing mice treated with vehicle or ENT starting on day 7 and tumors harvested for flow cytometry analysis on indicated days, representative plots shown. **A.** TIM3 versus Ki67 staining of tumor-infiltrating CD8 T cells on day 17. **B.** Number of tumor-infiltrating CD8_+_ T cells normalized to tumor mass on day 7 (no treatment), day 10, and day 17. **C.** TIM3 versus PD1 staining of tumor-infiltrating CD8_+_ T cells on day 17. **D.** Number of tumor-infiltrating CD8_+_ T cells normalized to tumor mass on day 7 (no treatment), day 10, and day 17. **E.** Expression of ICOS versus CD127 in TIM3_+_PD1_+_ (top) and TIM3_-_PD1_+_ (bottom) tumor infiltrating CD8_+_ T cells. **F.** Quantification of CD127 and ICOS staining by MFI in TIM3_+_PD1_+_ (top) and TIM3_-_PD1_+_ (bottom) tumor infiltrating CD8_+_ T cells, n=6 per cohort per time point. **G-J.** CD8_+_ T cells from tumor and draining lymph node of ENT-treated mice were sorted by TIM3 versus PD1 expression and labeled with CTV. Sorted cells were stimulated in culture for 108 hours with plate-bound anti-CD3/CD28 and IL-2. PD1 and TIM3 phenotype, CTV dilution, and quantification are shown for indicated CD8 populations sorted from the draining lymph node **(G-H)** or tumor mass **(I-J)**. Sorted cells from n=14 mice and cultured in duplicate, mean and SEM shown. Statistical significance calculated using t-test with Welch correction. **K-Q.** Transfer experiment of sorted CD8 progenitor and progeny. **K.** CD8+ T cells from tumor and draining lymph node of ENT-treated donor mice (CD45.2) were harvested on day 17, sorted by PD1+ and TIM3 expression, and transferred into recipient TS/A tumor-bearing mice (CD45.2). Tumor and draining lymph node of recipient mice harvested 10 days post transfer. **L-Q.** CD45.2+ CD8 T cells recovered from the tumor or lymph node of recipient mice. Concatenated plots showing PD1 versus TIM3 expression of recovered CD45.2 CD8 T cells from draining lymph node **(L)** or tumor **(O)** after transfer of TIM3+ (top) or TIM3- (bottom) cells. Quantification of PD1+, TIM3+ and PD1+, TIM3- CD8 T cells recovered from draining lymph node **(M)** or tumor **(P)** as a function of transferred phenotype. Slamf6 MFI for PD1+, TIM3+ and PD1+, TIM3- CD8 T cells recovered from draining lymph node **(N)** or tumor **(O)** as a function of transferred phenotype. Data show analysis of n=3 recipient mice per cohort and is representative of two independent experiments, mean and SEM shown. Statistical significance calculated using t-test. *p<0.05, **p<0.005, ^#^p<0.0005.

To directly test the proliferative capacity of TIM3^-^PD1^+^ and TIM3^+^PD1^+^ cells and determine whether the TIM3^-^PD1^+^ cells could give rise to TIM3^+^PD1^+^ cells, we sorted TIM3^-^PD1^+^ and TIM3^+^PD1^+^ cells from the draining lymph nodes (LNs) and tumors of ENT-treated mice, labeled them with CTV and cultured them with plate-bound anti-CD3, anti-CD28 and IL-2 for 4 days. We found that TIM3^-^PD1^+^ cells from the LNs gave rise to both TIM3^+^ and TIM3^-^ cells, whereas TIM3^+^PD1^+^ cells gave rise to predominantly TIM3^+^ progeny (**Figure 4G**). Although both populations proliferated in culture, the TIM3^-^PD1^+^ cells proliferated much more extensively (**Figure 4H**). We also sorted TIM3^-^PD1^+^ and TIM3^+^PD1^+^ cells from tumors and performed the same experiment. Again, we found that TIM3^-^PD1^+^ cells give rise to both TIM3^+^ and TIM3^-^progeny, whereas the TIM3^+^PD1^+^ cells gave rise to predominantly TIM3^+^ cells (**Figure 4I**). Moreover, the TIM3^-^PD1^+^ cells proliferated substantially more than the TIM3^+^PD1^+^ cells (**Figure 4J**). These results are consistent with the idea that the TIM3^-^PD1^+^ cells in cluster 5 are the precursors of the TIM3^+^PD1^+^ cells in clusters 1 and 2. To test this idea in vivo, ENT-induced TIM3+ PD1+ and TIM3- PD1+ CD8 T cells were sorted from CD45.2 tumor-bearing donors and transferred into CD45.1 tumor bearing recipient mice. Tumors and draining lymph nodes from recipient mice were then harvested 10 days following transfer and donor CD8 T cells analyzed by flow cytometry (**Figure 4K**). Transfer of TIM3- PD1+ cells led to greater recovery of donor CD8 T cells in the draining lymph node (**Figure 4L-M**) and tumor (**Figure 4O-P**). This difference was most pronounced in numbers of recovered TIM3-PD1+ progenitors, as these were not detected in the draining lymph node and were much less frequent in the tumor. Expression of slamf6, a surface marker faithfully correlating with TCF1 expression, was significantly higher in CD8 T cells recovered from TIM3- PD1+ transfer in both the draining lymph node (**Figure 4N**) and tumor (**Figure 4Q**). These data indicate that TIM3- PD1+ cells can self-renew and give rise to TIM3+ PD1+ cells but not the converse.

To test whether the accumulation of effector cells in the tumors of ENT-treated mice was due to local expansion of tumor-reactive T cells or to expansion in the LN and subsequent trafficking to the tumor, we treated tumor-bearing mice with combinations of ENT and the S1PR inhibitor, FTY720, beginning on day 7, and enumerated TIM3^-^Ki67^+^ and TIM3^+^Ki67^+^ CD8^+^ T cells in the LN and tumor on day 17. We found that ENT increased the TIM3^-^Ki67^+^ progenitor population and the TIM3^+^Ki67^+^ effector population in the LN (**Figure 5A-B**). However, the addition of FTY720 stymied the expansion of both populations (**Figure 5A-B**). In the tumor, ENT did not expand the TIM3^-^Ki67^+^ progenitor population, but dramatically increased the TIM3^+^Ki67^+^ effector population (**Figure 5C-D**). The addition of FTY720 abrogated the effects of ENT, but also dramatically reduced both the TIM3^-^Ki67^+^ and the TIM3^+^Ki67^+^ cells in the absence of ENT (**Figure 5C-D**), suggesting that the TIM3^-^Ki67^+^ progenitor cells in the tumor are coming from the LN and expanding in situ. The enhanced functional capacity of CD8^+^ T cells from ENT-treated tumors was also reversed by FTY720 (**Figure 5E**). As a result, treatment with FTY720 increased tumor growth (**Figure 5F**). Importantly, differences in CD8^+^ T cell frequencies within the tumor were not due to selective apoptosis incurred by ENT treatment (**Figure 5G-J**).

**Figure 5.**
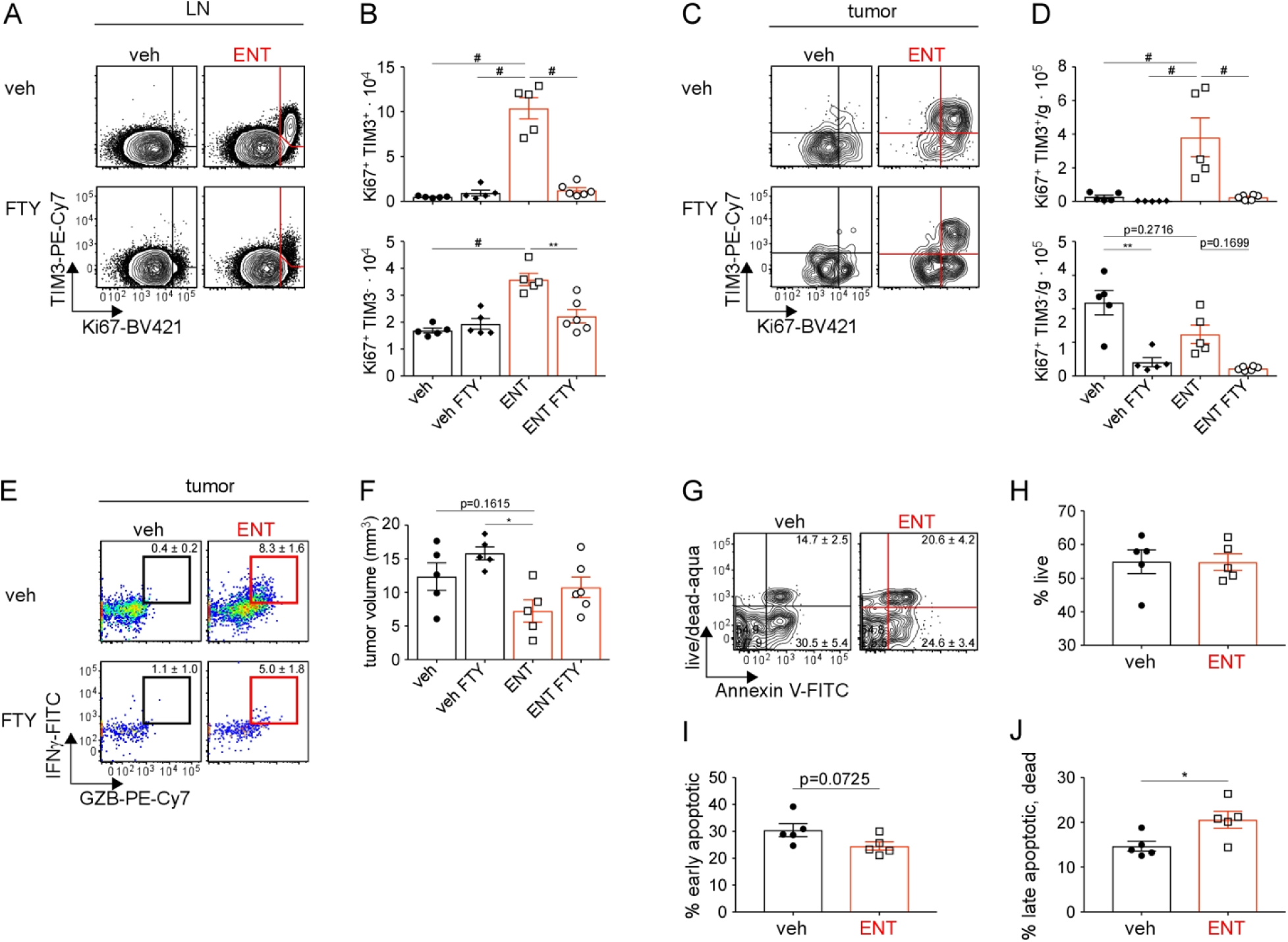
ENT-induced intratumoral progenitor and effector CD8 T cell populations are maintained by continuous trafficking. A-F. FTY720 given by IP injection to block S1PR-mediated trafficking during ENT treatment. ENT or vehicle daily treatment given on days 7-17. FTY720 or vehicle given on days 0, 1, 3, 5, 7, and 9. **A.** Tumor-draining lymph node (inguinal) harvested on day 17 and flow plots of CD8_+_ T cell TIM3 versus Ki67 staining shown. **B.** Enumerated counts of Ki67_+_ TIM3-positive and –negative CD8 T cells. Representative plots shown. **C.** Tumors harvested on day 17 and flow plots of intratumoral CD8_+_ T cell TIM3 versus Ki67 staining shown. **D.** Number of Ki67_+_ TIM3-positive and –negative CD8_+_ T cells normalized to tumor mass. Representative plots shown. **E.** Tumor harvested on day 17 and CD8_+_ T cells were restimulated ex vivo and stained for IFN_γ_ and GZB. Representative plots shown. **F.** Tumor mass on day 17 following ENT and/or FTY treatment described above, n=5-6 per cohort. Statistical significance determined using one-way ANOVA and Tukey’s method for multiple comparisons. **G-H.** TS/A tumor-bearing mice were treated daily with vehicle or ENT starting on day 7 and tumors harvested on day 17. **G.** Representative flow plots of CD8 T cells stained with live/dead dye and Annexin V. Corresponding frequencies of live **(H)**, early apoptotic **(I)**, and late apoptotic/dead cells **(J)** are shown, n=5 per cohort. Statistical significance determined by independent t-test. Mean and SEM shown. *p<0.05, **p<0.005, _#_p<0.0005.

### ENT treatment prevents chromatin restructuring to preserve proliferative capacity

Because HDAC inhibitors participate in large chromatin remodeling complexes, we asked if ENT treatment might impact chromatin reorganization to preserve proliferation and effector function (when given after activation). To address this, TS/A tumor-bearing mice were treated with ENT or vehicle beginning on day 7. Tumors were harvested on day 7 (baseline prior to treatment) or day 17. To prevent biasing of results by transient naïve T cells, activated tumor-infiltrating CD8 T cells were sorted by PD1 expression (positive) for ATAC-sequencing. Analysis demonstrated that PD1^+^ CD8 T cells in each cohort clustered separately on unsupervised PCA (**Figure 6A**). Comparison of differentially accessible regions (DARs) between day 17 vehicle- and ENT-treated revealed diffuse chromatin restructuring. 1045 total DARs were discovered, including 241 open and 804 closed (**Figure 6B**).

**Figure 6.**
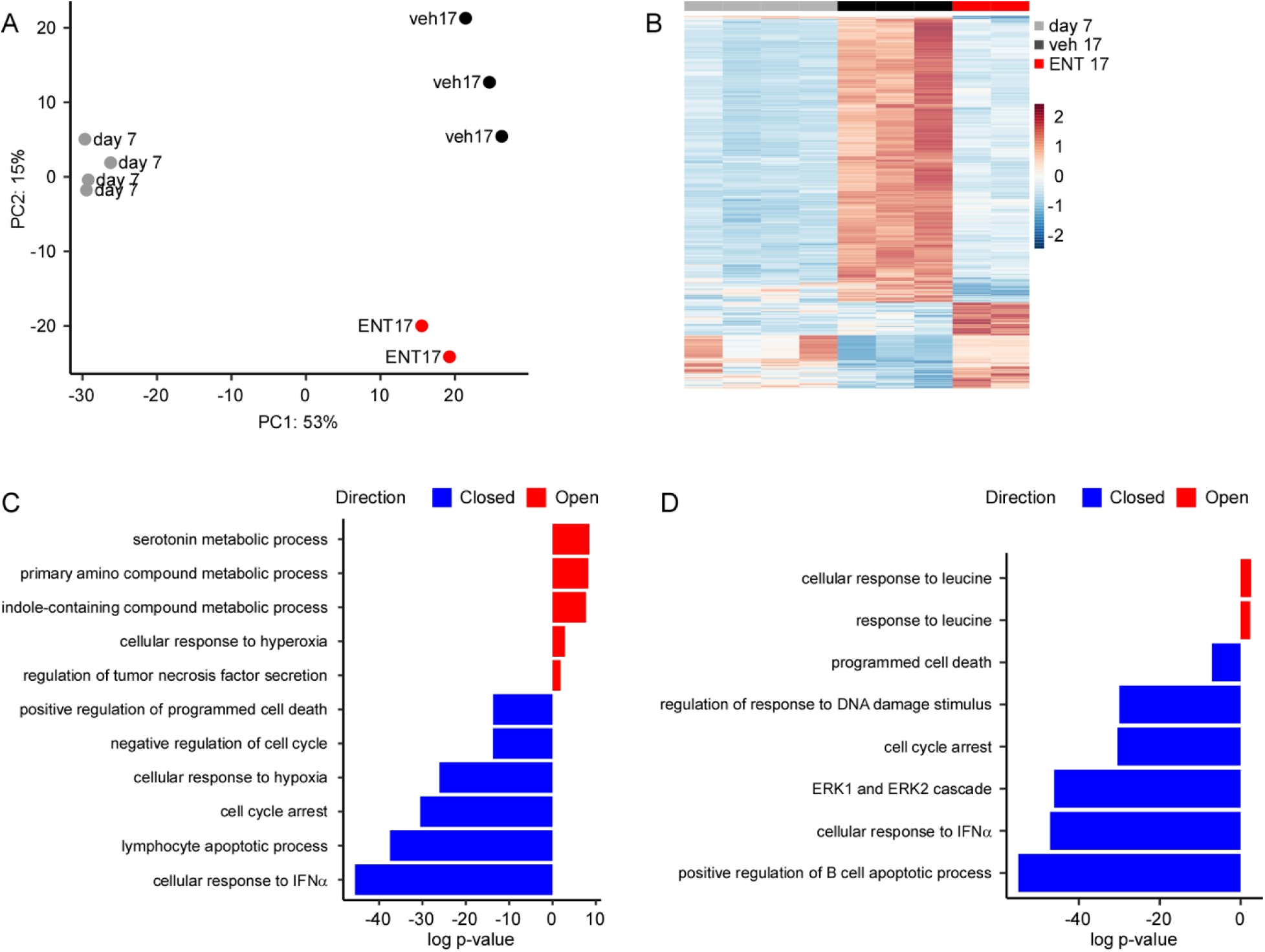
ENT treatment prevents opening of loci associated with cell cycle arrest, apoptosis, and exhaustion. TS/A tumor-bearing mice were treated daily with vehicle or ENT starting on day 7. Tumors were harvested on day 7 prior to treatment initiation (baseline) or on day 17 following ten days of vehicle or ENT. Activated infiltrating CD8 T cells were sorted and subjected to ATAC-sequencing. **A.** PCA plot of day 7, day 17 vehicle, and day 17 ENT samples. **B.** Heatmap of DARs when comparing day 17 vehicle versus day 17 ENT. **C.** Gene ontogeny identified by GREAT from comparison in B. Red corresponds to more open loci and blue to closed loci in day 17 ENT relative to day 17 vehicle. **D.** Gene ontogeny identified by GREAT using shared DARs between comparisons of day 17 vehicle versus day 7 and day 17 vehicle versus day 17 ENT. Red corresponds to open shared pathways and blue to closed shared pathways in day 7 and day 17 ENT relative to day 17 vehicle.

Comparing day 7 and day 17 vehicle samples, 4847 total DARs were discovered, including 612 open and 4235 closed (**Supplemental Figure 6A**). Numerous peaks that remained closed in day 7 and day 17 ENT samples became more accessible in day 17 vehicle samples, suggesting the most open chromatin belonged to the most dysfunctional CD8 T cells. Next, total DARs comparing day 17 ENT- and vehicle-treated were fed to the Genomic Regions Enrichment of Annotations Tool (GREAT) for gene ontogeny analysis. Less accessible pathways in ENT-treated CD8 T cells included negative regulation of cell cycling, cell cycle arrest, lymphocyte apoptotic process, and cellular response to IFNα (**Figure 6C**). When comparing day 7 versus day 17 vehicle samples, H3K27 methylation, H3K27 trimethylation, and DNA methylation were more accessible pathways in day 7 (**Supplemental Figure 6B**). Importantly, these pathways are commonly associated with more repressed chromatin. Specifically, trimethylation of H3K27 in CD8 T cells is highest in naïve and central memory subsets (relative to effector) [33] and correlates with increased memory formation and increased antigen-specific T cells on re-challenge [34].

Next, we compared day 17 vehicle versus day 7 and separately compared day 17 vehicle versus day 17 ENT samples. 612 upregulated DARs were in the former and 241 upregulated DARs were identified in the latter comparison, of which 98 DARs were shared. Among downregulated DARs, 4235 were identified when comparing day 17 vehicle versus day 7 and 804 when comparing day 17 vehicle versus day 17 ENT. Interestingly, nearly all closed DARs (746 of 804) discovered in the latter comparison were also present in the former (**Supplemental Figure 6C**). When shared up (98) and down (746) regulated DARs were fed to GREAT for gene ontogeny analysis, increased accessibility of pathways associated with programmed cell death, cell cycle arrest, ERK1/2 cascade, and cellular response to IFNα were found in day 17 vehicle samples relative to day 7 and day 17 ENT (**Figure 6D**). These data suggest ENT treatment keeps specific loci closed in tumor infiltrating CD8 T cells over time, loci associated with loss of proliferation, apoptosis, and exhaustion. Notably, sustained IFNα signaling [22, 24, 27] and the MAPK/ERK pathway [35] can drive CD8 T cell exhaustion. Inspection of canonical CD8 activation/effector loci revealed some differences in accessibility of *havcr2*, *pdcd1*, and *gzmb*; however, these did not reach statistical significance (**Supplemental Figure 6D**).

“Jump-starting” the T cell response restores the efficacy of ENT in established tumors

Our data show that ENT no longer enhanced T cell proliferation or function once they became exhausted (**Figure 1F**), suggesting that to treat established tumors, we would have to “jump-start” the T cell response. To test this possibility, we injected established tumors with an oncolytic HSV on days 13 and 15, treated mice daily with ENT for 2 weeks starting on day 13 or 19, and administered anti-PD1 starting on day 13 or 19 (**Figure 7A**). Control mice received ENT and anti-PD1 starting on day 19 (no HSV) or received HSV on days 13 and 15 followed by anti-PD1 on day 19 (no ENT). We found that the combination of ENT and anti-PD1 as well as the combination of HSV and anti-PD1 had no effect on tumor growth (**Figure 7B**), while HSV and ENT afforded moderate tumor control (**Supplemental Figure 7A**). However, the timed combination of HSV, ENT and anti-PD1 led to dramatic tumor regressions and even rejection, as did the simultaneous combination of those agents (**Figure 7B**), albeit with different kinetics.

**Figure 7.**
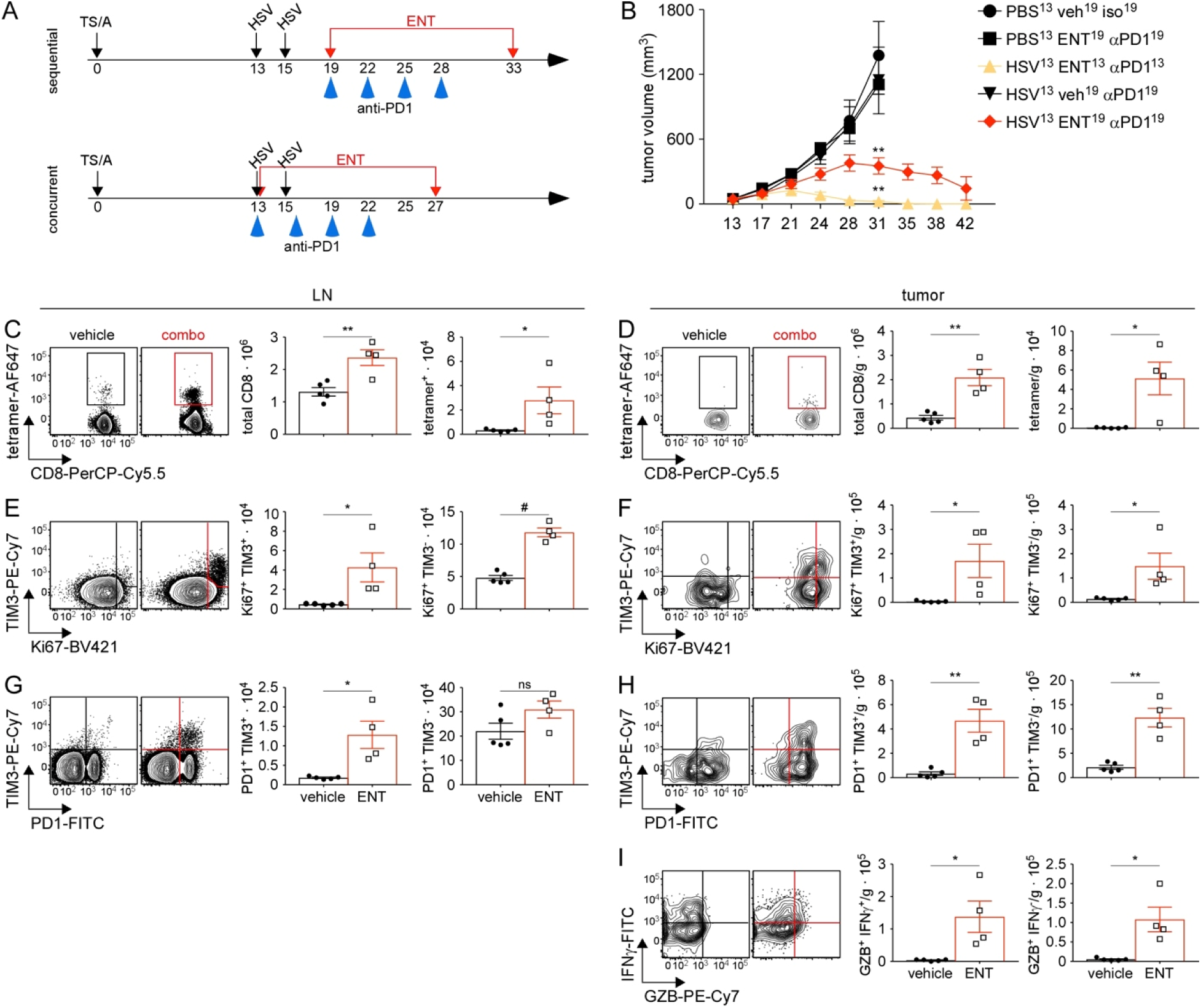
“Jump starting” the T cell response with an oncolytic HSV enables precise timing of HDAC inhibition and drives pronounced responses in late-stage tumors. **A.** Experimental design for “jump starting” the T cell response. TS/A tumors allowed to grow for 13 days prior to treatment, then HSV injected intratumorally on day 13 and 15. Daily ENT (red lines) and 200μg anti-PD1 (blue arrow heads) given starting on day 19. In the concurrent cohort HSV, ENT, and anti-PD1 started on day 13. **B.** Tumor growth curves following this strategy, n=3 per cohort. Experiment representative of second independent experiment where anti-PD1 started on day 24, n=4 per cohort. Statistical significance on day 31 determined by one-way ANOVA with Tukey’s method for multiple comparisons. **C-I.** Sequential experimental strategy was repeated with control (PBS, vehicle, isotype antibody) and combination (HSV then ENT and anti-PD1 starting on day 19) cohorts and CD8 T cell phenotype and function assessed in the draining lymph node and tumor (counts normalized to tumor mass) on day 28 by flow cytometry. Tumor-specific CD8^+^ T cells measured by tetramer staining. Total tetramer-positive and total CD8^+^ T cells in draining lymph node (**C**) and tumor (**D**). TIM3 versus Ki67 staining of CD8^+^ T cells and quantification in lymph node (**E**) and tumor (**F**). TIM3 versus PD1 staining of CD8^+^ T cells and quantification in lymph node (**G**) and tumor (**H**). CD8^+^ T cells from harvested tumor restimulated ex vivo and production of IFNγ and GZB measured by flow and quantified (**I**). n=4- 5 per cohort. Mean and SEM shown. Statistical significance calculated by t-test. *p<0.05, **p<0.005, ^#^p<0.0005.

To determine whether the efficacy of triple combination treatment (HSV^13^ENT^19^anti-PD1^19^) was mediated by a reinvigorated T cell response, we harvested tumors and LNs at day 28 from control and treated mice and analyzed CD8^+^ T cells. We found more total CD8^+^ T cells and tumor-specific CD8^+^ T cells in both the LNs and tumors of treated mice (**Figure 7C-D**). We also found more TIM3^-^Ki67^+^ progenitor-like cells and TIM3^+^Ki67^+^ effector cells in both the LNs and tumors of treated mice (**Figure 7E-F**). Similarly, the TIM3^-^PD1^+^ cells and the TIM3^+^PD1^+^ populations were also expanded in the LNs and tumors of treated mice (**Figure 7G-H**). Finally, we observed more CD8^+^ T cells making IFNγ and GZMB in the tumors of treated mice (**Figure 7I**). Taken together, these data suggest that “jump-starting” the T cell response with immune stimulatory agents like HSV generates a population of T cells that is susceptible to immune modulatory agents like ENT and anti-PD1, thereby enhancing anti-tumor immunity.

## DISCUSSION

Our data show that the timing of ENT administration relative to the state of T cell activation determines the functional outcome of anti-tumor immune responses. Given too early, ENT prevents T cell activation; given too late, it fails to reactivate exhausted T cells; but given at the peak of T cell activation, ENT sustains T cell proliferation and the differentiation of polyfunctional effector T cells. The narrow window during which ENT enhances T cell responses corresponds to a time when T cells have acutely activated gene expression programs controlling proliferation and effector functions [19]. As a result, ENT expands a population of proliferating PD1^+^TIM3^-^ progenitor-like T cells, which give rise to PD1^+^TIM3^+^ effector T cells that retain their functional capacity for prolonged periods. For established tumors with dysfunctional T cells, the T cell response must be “jump-started” (by an oncolytic virus in our experiments), in order for ENT and anti-PD1 to successfully enhance anti-tumor immunity. These data explain the poor clinical outcomes of two Phase 1b/2 clinical trials (NCT02915523, NCT02708680), which combined ENT with PD1 blockade, but failed to jump-start the T cell response.

The apparent disconnect between the preclinical data suggesting that ENT and anti-PD1 should be synergistic [15,16] and the disappointing clinical results [17], can be explained by how mouse experiments and clinical trials are performed. For implanted tumors, the act of injection initiates an acute immune response, in which T cells initially expand for about a week and then contract, with the remaining T cells progressively becoming more dysfunctional. Studied drugs can then be administered at a known time, often beginning the first week after implantation, which corresponds to the peak of the T cell response. However, this approach does not recapitulate what happens in patients, who often arrive in the clinic months or years after their tumors first begin to grow. Consequently, tumor-specific T cells in these patients are likely dysfunctional [36], and fail to respond to the combination of ENT and anti-PD1. To overcome this unresponsiveness, the T cell response must be acutely re-initiated in a way that makes it susceptible to the effects of ENT and PD1 blockade. To accomplish this goal, we used an oncolytic HSV to kill tumor cells, release tumor antigens and cause inflammation – conditions that initiate primary immune responses or restimulate resting memory T cells [37,38]. Given that the simultaneous administration of HSV, ENT and anti-PD1 successfully enhanced the CD8^+^ T cell response and effectively eliminated tumors, we conclude that HSV most likely reawakened resting memory T cells, while possibly also priming naïve T cells. This scenario is similar to one in which the combination of anti-PD1 and TIM3 blockade reawakened virus-specific CD8^+^ T cells 200 days after infection [39].

Mechanistically, our results show that, when administered at the proper time, ENT treatment expands and sustains a proliferating progenitor-like population of T cells. Functionally similar progenitor-like or stem-like CD8 T cells are observed in the context of chronic virus infection and can be identified based on the expression of CXCR5 and TCF1 [40,41]. Although the progenitor cells we describe express markers like CD127, ICOS, and TCF1 (at the protein level) that correspond to central memory or progenitor-like T cells, they do not express CXCR5. This difference may reflect the way T cells respond to viruses or tumors or may imply that ENT enables a distinct class of progenitor cells. Nevertheless, our findings are consistent with reports identifying PD1^+^TIM3^-^ progenitor-like cells that give rise to PD1^+^TIM3^+^ effector T cells [40–42]. Thus, appropriately timed HDAC inhibition can therapeutically induce and maintain a population of CD8^+^ T cells that responds positively to PD1 blockade, thereby sensitizing formerly non-responsive tumors to checkpoint inhibitors.

In sum, our data demonstrate that the impact of class I HDAC inhibition on anti-tumor immunity is entirely dependent upon the timing of inhibition relative to T cell activation. Appropriately timed HDAC inhibition therapeutically expands progenitor CD8^+^ T cells that give rise to robust effector cells that retain cytotoxic functions for prolonged periods. Inappropriately timed HDAC inhibition however, fails to activate T cells or prevents their activation, likely explaining their poor clinical performance against solid tumors. To overcome this limitation and empower productive anti-tumor immunity, we propose that the T cell response must be jump-started immediately prior to HDAC inhibition by oncolytic agents like HSV or by other agents that cause the immunogenic cell death of tumor cells [43].

## METHODS

### Mice, tumor administration, and treatment regimens

BALB/c mice were purchased from Charles River Laboratories International or Jackson Laboratory. C57BL/6J mice were purchased from the Jackson Laboratory and bred in the University of Alabama at Birmingham vivarium. All mice were used between 7-8 weeks of age. Mice were injected with 1 x 10^5^ TS/A cells in the second mammary pad or 8 x 10^4^ MC38-OVA cells in the flank on day 0. Tumor volume was calculated by measuring tumor length (L) and width (W) using calipers and using the formula: volume = 0.4 x L x W^2^. Tumor bearing mice were treated with 20 mg/kg ENT daily for two weeks starting at the indicated times. 200μg of anti-PD1 (BioXCell, clone RMP1-14) or isotype control antibody (BioXCell, clone 2A3) were administered by intraperitoneal injection on the days indicated. The IL-12-expressing HSV, M002 [44], was diluted to 3.33 x 10^8^ PFU/mL in sterile PBS and 30 μL, or 1 x 10^7^ PFU, was injected into the tumor mass.

### Cell culture

The TS/A murine mammary adenocarcinoma was cultured in DMEM supplemented with 10% fetal bovine serum (FBS; both from Hyclone Laboratories, Inc.) and 2% L-glutamine, penicillin, streptomycin solution (Sigma) as described [15]. MC38 cells were purchased from ATCC and stably transfected with an ovalbumin (OVA) expression plasmid (pHb-Apr-1-OVA described by Pulaski et al. [45]). MC38-OVA cells were cultured in RPMI 1640 supplemented with 5% FBS; 2% L-glutamine, penicillin, streptomycin solution; and 0.1% gentamicin (Gibco), passaged by dissociation with 0.05% trypsin, 0.53 mM EDTA (Corning), and transferred to fresh media. Cell lines were thawed and passaged at least 3 times prior to use and not used after 7 passages.

### Chemicals and reagents

Entinostat (LC laboratories) stocks were prepared in DMSO, then stock solutions diluted to the final concentration in a vehicle of 15% DMSO, 20% Kolliphor EL, and 65% PBS, 0.2um filtered. The S1PR inhibitor FTY720 (Fingolimod; Cayman Chemical) was prepared at 2 mg/mL in ethanol and diluted to 0.2 μg/μL in PBS prior to intraperitoneal injection of 40 μg total FTY720 per injection as described.

### Tumor dissociation and T cell restimulation ex vivo

Tumors were excised with a scalpel, weighed using an AL54 analytical balance (Mettler Toledo), and minced with scissors. Tumor fragments were incubated in 2 mL of RPMI-1640 supplemented with 5% FBS, 1.25 mg collagenase (c7657, Sigma) and 150 U DNase (d5025, Sigma) under constant agitation at 200 RPM for 35 minutes at 37°C. Dissociated tumors were mashed through 70 μm nylon cell strainers (Corning), using a 5 ml syringe plunger. T cells were restimulated ex vivo by culture in RPMI-1640 supplemented with 5% FBS, 5 ng/mL phorbol 12-myristate 13-acetate (Sigma), 65 ng/mL ionomycin (ThermoFisher Scientific), and 10 μg/mL brefeldin A (Sigma) for 4-5 hours at 37°C. Cells were washed in PBS with 2 μg/mL brefeldin A.

### Flow cytometry, antibodies, and T cell sorting

For immunophenotyping, single cell suspensions were washed and resuspended in staining media (PBS with 2% donor calf serum and 10 μg/mL FcBlock (2.4G2, BioXCell)). Staining for surface proteins was performed for 30 minutes at 4°C. Fixation prior to intracellular staining was done using the eBioscience FoxP3 Fixation Kit (ThermoFisher Scientific) for 30 minutes at room temperature or using the BD Fixation/Permeabilization Solution Kit (BD Biosciences) for 20 minutes at 4°C. Intracellular staining was performed on fixed cells permeabilized as described by the manufacturer for at least 45 minutes at 4°C. All staining was performed in the dark. Antibodies against CD4 (GK1.5), Tbet (4B10), and Ki67 (16A8) were purchased from BioLegend. Antibodies against IFNγ (XMG1.2), TNF (MP6-XT22), and CD8 (53-6.7) as well as the FITC Annexin V kit were purchased from BD Biosciences. Antibodies against CD3 (17A2), CD45.2 (104), CD127 (A7R34), TIM3 (RMT3-23), PD1 (J43), ICOS (C398.4A), and granzyme B (NGZB) were purchased from eBioscience. Live/dead fixable dyes were obtained from Life Technologies. Cell trace violet and anti-Id2 (ILCID2) were purchased from Invitrogen. Anti-TCF1 (812145) was purchased from R&D Systems. Tetramers containing murine leukemia virus env peptide (SPSYVYHQF) were acquired from the NIH Tetramer Core Facility. Samples were run on a BDFACS Canto II system (BD Biosciences) and data analyzed using FloJo version 9.9. A typical gating strategy is shown in **Supplemental Figure 1**.

For T cell sorting, dead cells were first removed from prepared single cell suspensions using the MACS Dead Cell Removal Kit (Miltenyi Biotec) per the manufacturer’s protocol. Bound bead-cell mixtures were passed over MACS LS columns (Miltenyi Biotec) and live cell containing eluates collected. Cell solutions were then stained for CD45.2, CD8 or CD4, live cells as described above and sorted using the BDFACS Aria II system (BD Biosciences). T cells were collected directly into RPMI-1640 supplemented with PBS (Corning) with 0.04% Bovine Serum Albumin (Thermo Fisher) for single cell analysis or Qiagen RLT buffer (Qiagen) plus 2-mercaptoethanol (Sigma) for bulk sequencing.

### Bulk RNA sequencing and analysis

Total RNA was purified using the Quick-RNA MicroPrep kit (Zymo Research) and used as input for the SMART-seq v4 cDNA synthesis kit (Takara). 200 pg of cDNA was used as input for the NexteraXT kit (Illumina) for tagmentation, PCR amplification and indexing. Final libraries were quantitated by Qubit, pooled at equimolar ratio, and sequenced on a NextSeq500 using 75bp SE chemistry at the UAB Helfin Genomics Core. FASTQ reads were mapped to the mm10 mouse genome using STAR [46] with the default parameters and the UCSC KnownGene transcriptome database [47]. The coverage at exons for all unique Entrez genes was computed using costom R/Bioconductor scripts and the GenomicRanges package [48]. The count matrix was processed using the R package DESeq2 (version 1.20.0) [49], and genes with total counts lower than 10 were excluded. For visualization purpose, the regularized-logarithm transformation was conducted using *rlog* function and principal component analysis (PCA) was performed on the transformed data. Normalized counts for genes associated with T cell function and proliferation were generated using *plotCounts* function. For differential expression analysis, *DESeq* function was run on the raw counts and three comparisons were conducted: day 10 ENT vs day 10 vehicle; day 17 ENT vs day 17 vehicle; and day 17 ENT vs day 10 ENT. Genes with an adjusted p value smaller than 0.1 and an absolute value of log2 fold change greater than 1 were defined as differentially expressed genes (DEGs). The DEGs obtained from the two ENT-vs-vehicle comparisons were further examined and classified into three subsets based on the time points when they were found differentially expressed. For GSEA, all detected genes were ranked by multiplying the -log_10_ of the P-value from DESeq2 by the direction (i.e., positive or negative) of the fold-change. The resulting ranked list was used as input for the GSEA PreRanked analysis. The RNA-seq data from this study is available from the NCBI Gene Expression Omnibus (GEO).

### Single cell RNA-seq and analysis

Single cell libraries were prepared according to the protocol for Chromium Single Cell 3’ Reagent Kit v2 (10X Genomics). Briefly, sorted CD8^+^ T cells, barcoded single cell 3 gel beads and partitioning oil were loaded onto a Single Cell A Chip, and single cell Gel Bead-In-EMulsions (GEMs) were generated by Chromium controller. Barcoded cDNA was then produced by the incubation of GEMs and amplified by PCR after the GEMs were broken to generate sufficient mass. Sequencing libraries were finally constructed with the P5 primer, sample index and P7 primer added, and sequenced by Illumina Nextseq machine at the UAB Helfin Genomics Core.

An aggregate count matrix was generated following the CellRanger pipeline (version 2.1.1) provided by 10X Genomics. Briefly, cell barcodes were demultiplexed, and reads were aligned to the mm10 genome and normalized across samples for aggregation. We next processed the gathered matrix using the R package Seurat (version 2.3.1) [50]. Cells with the total number of genes between 200 and 5000 and UMI lower than 60,000 were kept. We further excluded cells whose mitochondrial gene counts accounted for more than 10% of total counts. 10569 cells with 15354 genes remained following the data preprocessing. Highly variable genes were detected using *FindVariableGenes* function in Seurat and genes whose log mean was between 0.0125 and 4.5 and dispersion was greater than 0.5 were selected for dimensionality reduction. PCA was first implemented and the top 14 significant PCs were then used for tSNE transformation and graph-based cluster analysis. By plotting the expression of Cd8a, Cd8b1 Cd3e, Cd247, Lck and Bst2, we found that 4 cell clusters (cluster 8, 12, 15 and 16) expressed high levels of Bst2 but not hallmark T cell genes such as Cd3e, Cd247 and Lck. This suggests that they represented non-T cell contaminants and were thus discarded from downstream analyses.

Following the exclusion of these clusters, we re-performed the normalization as suggested by the Seurat package documentation and detected 1937 highly variable genes from the cleaned raw count. We then implemented PCA, tSNE and graph-based clustering as described above and obtained 13 clusters. The cluster assignment of cells from ENT- or vehicle-treated animals was compared to reveal the effects of the treatment. To determine the characteristics of a cell cluster, gene expression in cells from that cluster was compared with those from other clusters using *FindMarkers* function. Both of the *min.pct* and *logfc.threshold* parameters were set to 0 so that no gene was filtered out. The 1937 variable genes were then ranked based on log fold change generated from this inter-cluster comparison. Consequently, 13 gene lists were generated, which contained the same genes but in different orders. Top genes from each ordered gene list were selected for visualization. To further associate the feature genes of cluster 1, 2, 5 and 10 with pathways, GSEA Preranked was performed with default parameters on the ranked gene lists corresponding to these 4 clusters.

For pseudotime analysis the data was imported to Monocle [51] using the *importCDS* function and the top variable genes were set using the *FindVariableGenes* function. Pseudotime ordering of cells was performed using the *orderCells* function using the DDRTree method.

### ATAC-sequencing and analysis

Tumor-infiltrating PD1^+^ CD8 T cells were sorted as described above. Cells were resuspended in 25 ml tagmentation buffer (1× TD Buffer (Illumina), 0.1 % Tween-20, 0.2% Digitonin, and 2.5 ml Tn5) and incubated at 37 °C for 1 hour, diluted 2 × in Lysis Buffer (300 mM NaCl, 100 mM EDTA, 0.6% SDS, 1.6 mg Proteinase-K), and incubated for 30 minutes at 40 °C. Size selection using SPRI-beads isolated low molecular weight DNA which was then PCR amplified using 2 × HiFi HotStart Ready mix (Roche) and Nextera Indexing Primers (Illumina). A SPRI-bead purification was performed, samples were quality checked for ATAC-seq specific patterning on a bioanalyzer and were pooled at an equimolar ratio and sequenced on a NextSeq550 using 75bp PE chemistry at the UAB Helfin Genomics core.

ATAC-seq data analysis was performed as previously described [52,53]. Briefly, FASTQ reads were mapped to the mm10 mouse genome using Bowtie2 [54] with the default settings. PCR duplicate reads were marked using PICARD (http://broadinstitute.github.io/picard/) and removed from subsequent analyses. MACS2 [55] was used for peak calling, and peaks from all samples were merged using HOMER’s mergePeaks function [56]. The GenomicRanges package was used to generate a raw and normalized count matrix from the mergePeaks file, which was then processed using the R package DESeq2. For visualization purpose, the regularized-logarithm transformation was conducted using *rlog* function and PCA was performed on the transformed data. For differential accessibility analysis, *DESeq* function was run on the raw counts, and loci with an adjusted p value smaller than 0.1 and an absolute value of log2 fold change greater than 1 were defined as differentially accessible regions (DARs). For gene ontology analysis, DARs were separated into open and closed regions, and their genomic ranges were generated using makeGRangesFromDataFrame function of the R package GenomicRanges. The genomic ranges were then used as input for Genomic Regions Enrichment of Annotations Tool (GREAT) [57].

### Statistical analysis

Statistical analysis was conducted in GraphPad Prism version 7.0a. Statistical significance between two group means at different time points was done using multiple unpaired independent t-tests, using the Holm-Sidak method to correct for multiple comparisons, and assuming alpha equal to 0.05. Two-way ANOVA with Tukey’s method for multiple comparisons was used for determination of significance between more than two group means. Error bars represent standard error. Specific tests used and sample numbers of each comparison are provided in corresponding figure legends.

## Data Availability

The bulk RNAseq and single cells RNAseq are archived under accession numbers GSE140285 and GSE140522, respectively. All other data are available upon request.

## Acknowledgements.

The authors would like to thank Uma Mudunuru and Scott Simpler for animal husbandry; Sagar Hanamanthu and Marion Spell of the Comprehensive Flow Cytometry Core for flow sorting expertise; and Antonio Di Stasi, Robin Lorenz, Tim Townes, and Casey Weaver for helping guide progression of this work.

## Author contributions

Tyler R. McCaw designed, performed and interpreted experiments and wrote the manuscript.

Mingyong Liu designed, performed, interpreted experiments, and conducted RNAseq analysis.

Christopher D Sharer conducted RNAseq analysis.

Jeremy M Boss conducted RNAseq analysis.

Tian Mi conducted RNAseq analysis.

Alexander F Rosenberg assisted in RNAseq analysis.

Mei Li performed experiments.

Rebecca C. Arend designed and interpreted experiments.

Andres Forero designed and interpreted experiments.

Donald J. Buchsbaum designed and interpreted experiments and edited the manuscript.

James M Markert designed and interpreted experiments and edited the manuscript.

Troy D. Randall designed and interpreted experiments and edited the manuscript.

