## Supplemental Figures 1-7 for "Jump-starting the T cell response in established tumors"

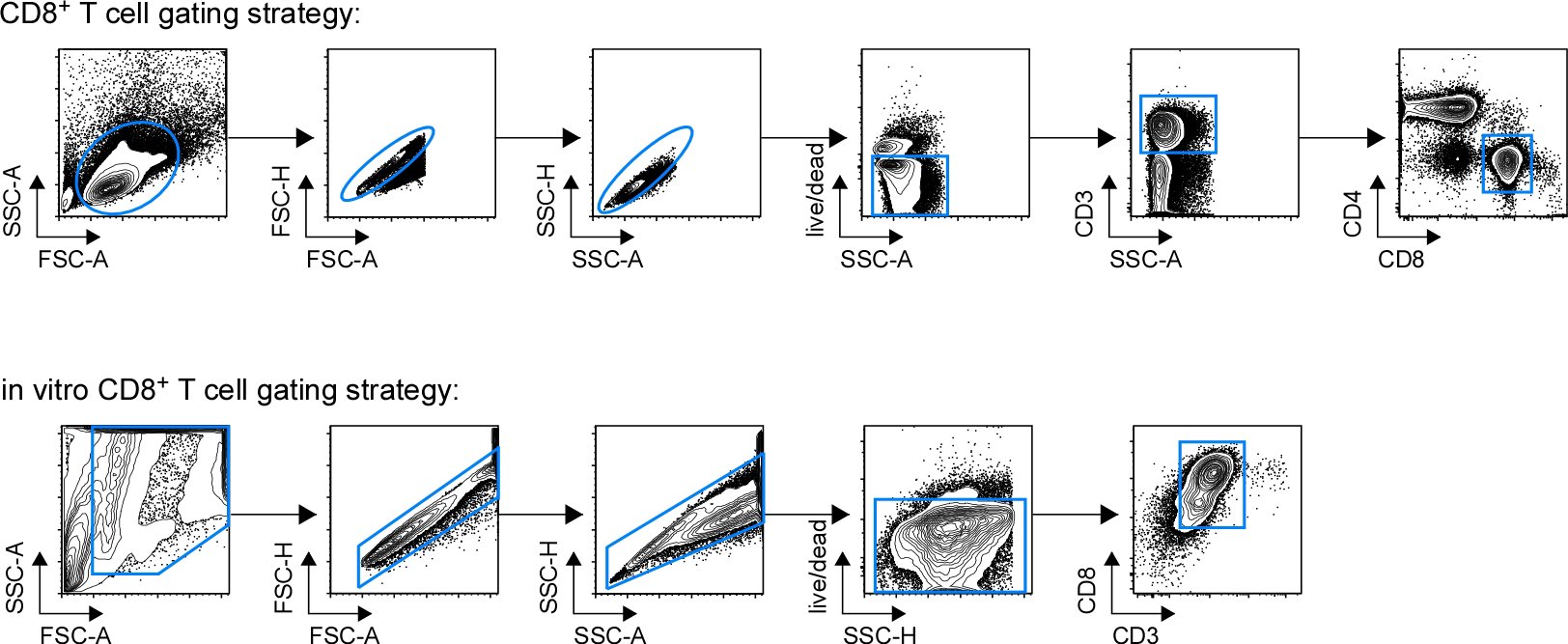

**Supplemental Figure 1:** Flow cytometry gating strategies. Multicolor flow cytometry gating strategies for CD8+ T cells *in vivo* (top) and *in vitro* (bottom).

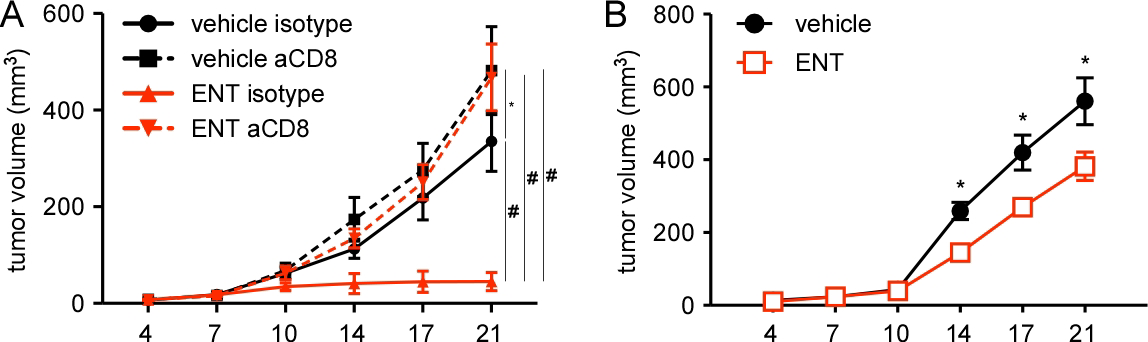
**Supplemental Figure 2:** CD8 T cells mediate the anti-tumor effects of ENT. **A.** Tumor growth of 8e4 MC38-ova cells injected subcutaneously into the flank of B6 wildtype mice and daily vehicle or ENT treatment plus 200mg isotype control or anti-CD8 antibody starting on day 7, n=5 per cohort. Statistical significance on day 21 determined by one-way ANOVA with Tukey’s method for multiple comparisons. **B.** Tumor growth of 8e4 MC38-ova cells injected subcutaneously into the flank of B6 TCR-beta KO mice and daily vehicle or ENT treatment started on day 7, n=5 per cohort. Statistical significance determined by independent t-tests at each time point. Mean and SEM shown, ^#^p < 0.0005.

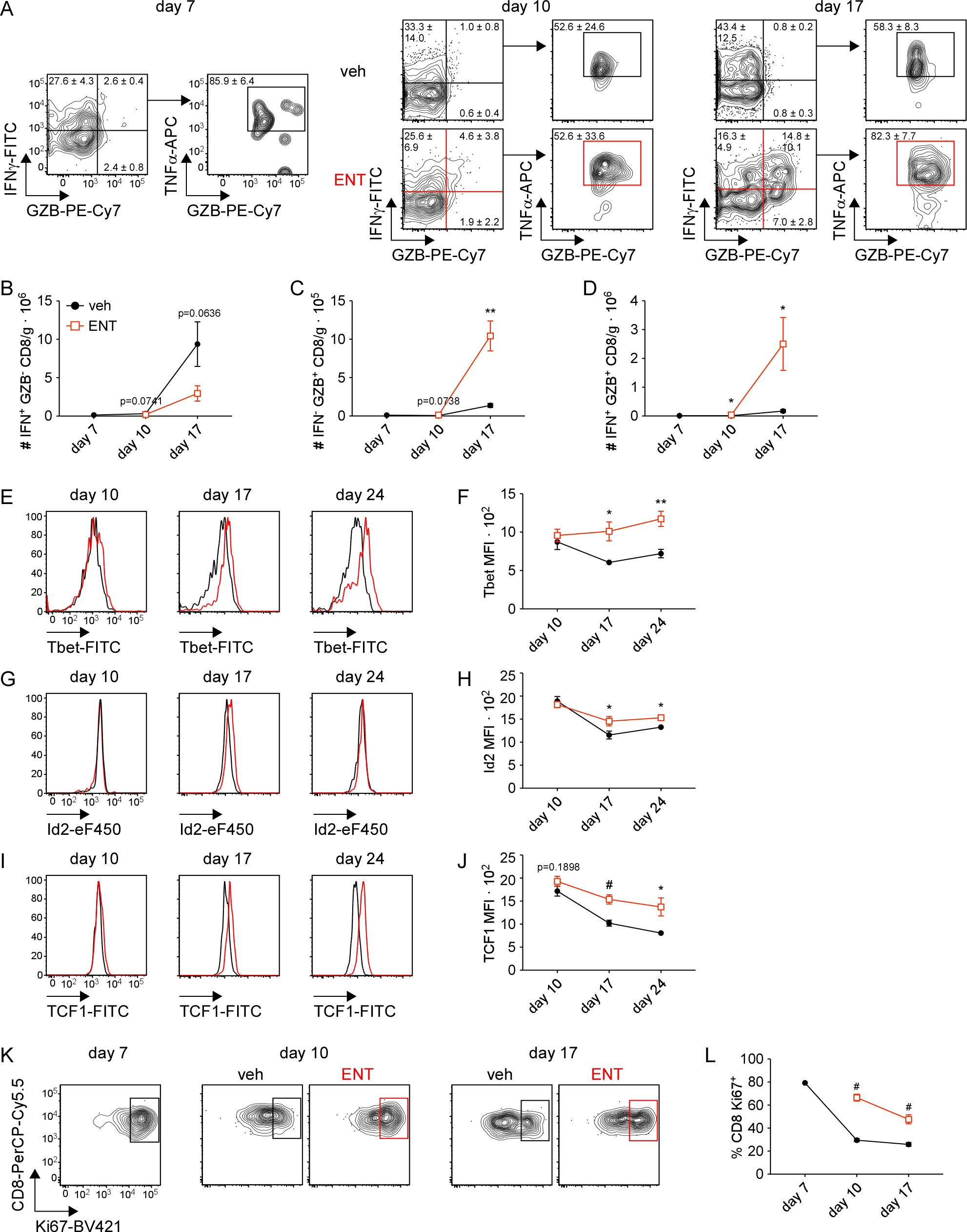

**Supplemental Figure 3:** Protein-level confirmation of identified DEGs in tumor-infiltrating CD8 T cells over time. **A-L.** TS/A tumor-bearing mice were treated with daily vehicle or ENT from day 7-20 (or until harvest), tumors harvested on indicated days, and T cell ex vivo restimulation done prior to intracellular cytokine staining for flow cytometric analysis, n=5-7 per cohort for 1-2 independent experiments per time point. Statistical significance determined by independent t-test at each time point. Mean and SEM shown. **A.** Representative flow plots of intratumoral CD8 T cell cytokine production on days 7 (no treatment), 10, and 17. Arrow to TNFα versus GZB plot shows TNFα production by IFNγ and GZB double-positive cells. Frequency mean ± standard deviation shown for each gate. **B.** Number of IFNγ single-positive CD8 T cells normalized to tumor mass. **C.** Number of GZB single-positive CD8 T cells normalized to tumor mass. **D.** Number of IFNγ, GZB double-positive CD8 T cells normalized to tumor mass. **E.** Representative flow plots of Tbet staining of intratumoral CD8 T cells. **F.** Frequencies of Tbet-positive intratumoral CD8 T cells. **G.** Representative flow plots of Id2 staining of intratumoral CD8 T cells. **H.** Frequencies of Id2-positive intratumoral CD8 T cells. **I.** Representative flow plots of TCF1 staining of intratumoral CD8 T cells. **J.** Frequencies of TCF1-positive intratumoral CD8 T cells. **K.** Representative flow plots of Ki67 staining of intratumoral CD8 T cells. **L.** Frequencies of Ki67-positive intratumoral CD8 T cells. *p<0.05, **p<0.005, ^#^p < 0.0005.

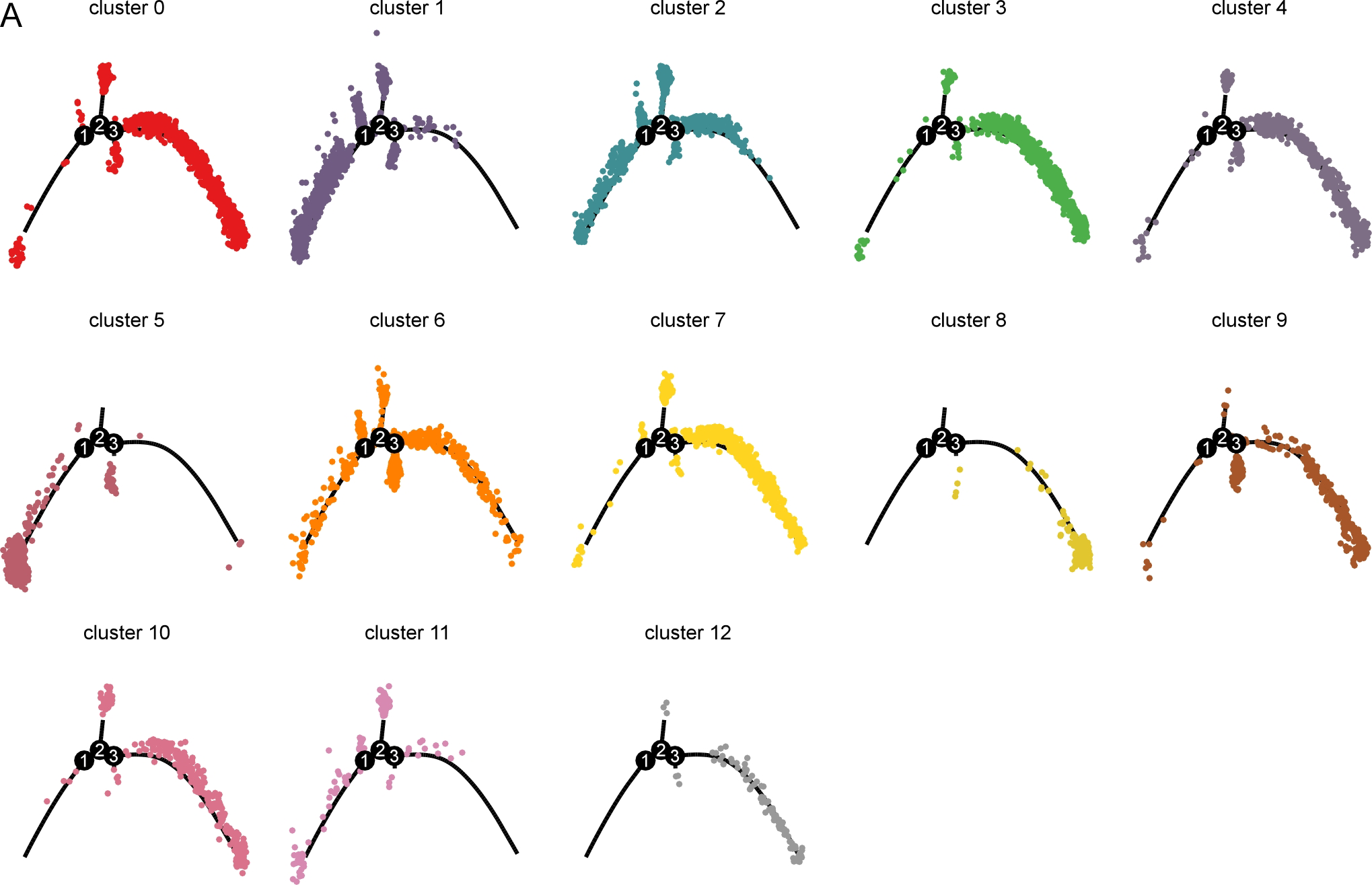

**Supplemental Figure 4:** Pseudotime trajectory analysis shows HDAC inhibitor treated clusters are less differentiated. **A.** TS/A tumor-bearing mice were treated with ENT or vehivle starting on day 7 and tumor-infiltrating CD8 T cells sorted on day 17 for single cell RNA-sequencing. Two independent experiments, n=2 per cohort per experiment. Trajectory plots with each individual cluster are overlaid. Number of cluster shown is provided above each plot.

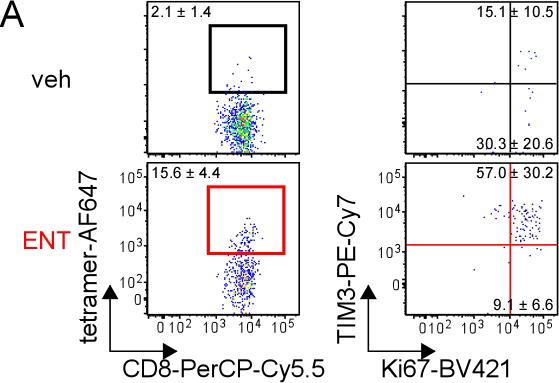

**Supplemental Figure 5:** Intratumoral tumor-specific CD8 T cells are expanded by ENT treatment. **A.** TS/A tumor-specific CD8 T cells within the TME on day 17. Ki67 versus TIM3 staining of tetramer-positive cells is shown. Representative plots with mean ± standard deviation, n=4-5 per cohort.

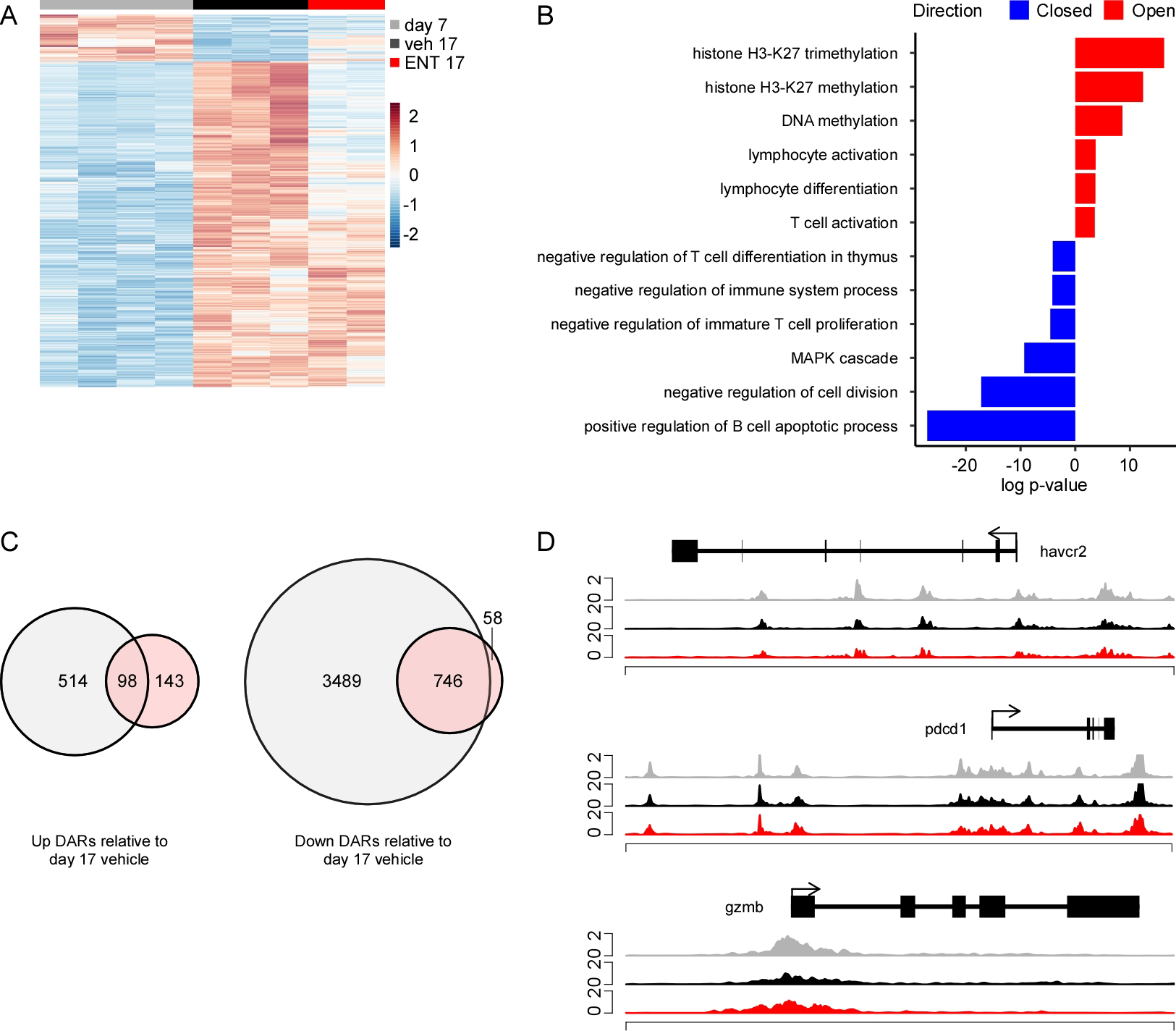

**Supplemental Figure 6:** ATAC-sequencing analysis of ENT treated CD8 T cells suggests different chromatin states and regulation. TS/A tumor-bearing mice were treated daily with vehicle or ENT starting on day 7. Tumors were harvested on day 7 prior to treatment initiation (baseline) or on day 17 following ten days of vehicle or ENT. Activated infiltrating CD8 T cells were sorted and subjected to ATAC-sequencing. **A.** Heatmap of DARs when comparing day 7 to day 17 vehicle. **B.** Gene ontogeny identified by GREAT using DARs from A. **C.** Venn diagrams showing total and shared DARs discovered when comparing day 17 vehicle versus day 7 and day 17 vehicle versus day 17 ENT. **D.** Accessibility at *havcr2*, *pdcd1*, and *gzmb* loci.

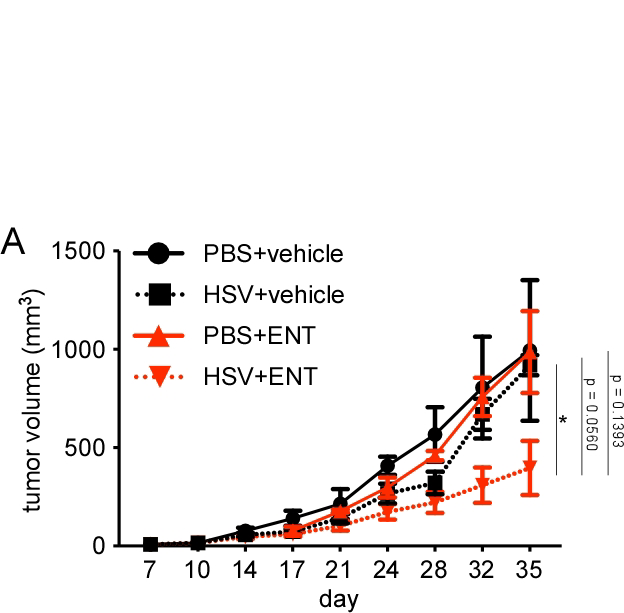

**Supplemental Figure 7:** The combination of HSV and ENT moderately impairs growth of established TS/A tumors. **A.** Balb/c mice were injected with TS/A cells on day 0 and tumors allowed to grow unencumbered until day 13. Oncolytic virus delivered by intratumoral injection on days 13 and 15 and ENT treatment started on day 19. The combination of HSV and ENT impaired growth of established tumors while neither agent was effective in isolation. n = 5 mice per cohort; data is representative of two independent experiments. Statistical significance at day 35 determined by t-test with Bonferroni correction for multiple comparisons. Mean and SEM shown. *p<0.05, **p<0.005, ^#^p<0.0005.
