## Supplemental Table 1 for "Jump-starting the T cell response in established tumors"

**Supplemental Table 1: Characteristic DEGs and nomenclature by cluster**

| **Cluster** | **Feature** | **Nomenclature** |
| --- | --- | --- |
| 0 | Klf2^hi^ Ly6c1^hi^ CD28^hi^ Tcf7^hi^ | Tcf7^hi^ resting |
| 1 | Ifng^hi^ Gzmb^hi^ Gzmd^hi^ Gzme^hi^ Irf8^hi^ Csf1^hi^ Pdcd1^+^ Havcr2^hi^ | Havcr2^hi^ effector |
| 2 | Gzma^hi^ Gzmb^+^ Gzmd^-^ Gzme^-^ Havcr2^int^ Prss2^hi^ | Gzma^+^ effector |
| 4 | Slamf6^hi^ Bcl2^+^ CD28^-^ | Slamf6^hi^ resting |
| 5 | Stmn^hi^ Tuba1b^hi^ Ly6e^hi^ Mki67^hi^ Cdk1^hi^ | Replicating |
| 6 | Lag3^hi^ Eomes^hi^ Tox^hi^ Pdcd1^+^ Cd160^+^ Dusp4^hi^ Eea1^hi^ | Dysfunctional effector |
| 7 | Fasl^+^ Eomes^+^ Gzmk^+^ Itga4^hi^ Gpr183^hi^ | Gzmk^+^ effector |
| 9 | Cd7^hi^ Cd83^+^ Klra5^hi^ Klra7^hi^ Ikzf2^hi^ | Helios^+^ resting |
| 10 | Isg15^hi^ Isg20^hi^ Ifit1^hi^ Ifit3^hi^ | ISG expressing |
| 12 | Ramp1^hi^ Sepp1^hi^ Hk2^hi^ | Ramp1^hi^ |
